# Structural and Functional Insights into Alkaline-pH Sensing by the Alka Channel

**DOI:** 10.1101/2025.10.29.685377

**Authors:** Tingwei Mi, Yuhang Wang, Yan Zhao, Yali V. Zhang

## Abstract

The ability to detect environmental pH is essential for survival. In *Drosophila melanogaster*, the taste receptor Alka mediates alkaline food sensing, but the structural and functional mechanisms of Alka-mediated high-pH sensation have remained unclear. Here, we report the cryo-electron microscopy structure of Alka at near-atomic resolution. Alka assembles into a homopentamer and undergoes conformational changes between neutral and alkaline pH conditions. At nearly neutral conditions, a lysine residue (K229) in the channel activation domain forms a stabilizing salt bridge, maintaining the channel in a closed state. Under alkaline conditions, deprotonation of the K229 side chain disrupts this interaction, resulting in channel opening. Functional studies in heterologous cells and intact flies show that mutation of K229 significantly impairs alkaline pH-dependent Alka activation, demonstrating its essential role in pH sensing. Our findings uncover a previously unrecognized structural mechanism underlying alkaline pH detection and identify a lysine residue as a molecular switch for alkaline pH-mediated channel gating. Moreover, our work establishes hydroxide ions as a distinct class of ion channel ligands and reveals a molecular innovation of chemosensory receptors by which animals broaden their range of chemical senses to detect and avoid strongly alkaline environments.

## Introduction

pH is fundamental to life, and animals rely on specialized chemical sensors to detect and avoid harmful extremes ^1–9^. Importantly, environmental pH is not constant — it can fluctuate dramatically and reach highly basic levels in ecological contexts such as alkaline lakes, soils, and decomposing organic matter ^10–13^. Thus, understanding how animals sense alkaline stimuli is essential for revealing how they adapt to chemically diverse environments ^14–16^. Until recently, however, the molecular basis of sensing highly alkaline conditions was unknown ^17^. We previously identified Alka, the first alkaline taste receptor, which enables *Drosophila* to detect strongly basic compounds in food ^6^. Alka belongs to the pentameric ligand-gated ion channel (pLGIC) superfamily, an evolutionarily ancient group found across both prokaryotes and eukaryotes ^18,19^. Unlike canonical pLGICs, which are activated by neurotransmitters such as glycine ^20,21^, gamma-aminobutyric acid (GABA) ^22,23^, and acetylcholine ^24–26^, Alka shows very low sequence identity to its relatives, lacks a conventional ligand-binding pocket, and is directly activated by extracellular alkaline pH ^6^. Alka therefore provides a unique model to investigate how ion channel structures are adapted to detect alkaline pH stimuli.

Determining its three-dimensional (3D) structure is crucial to uncovering the molecular basis of this unusual pH-gating mechanism and, more broadly, to understanding how sensory systems evolve to detect ecologically relevant cues outside physiological ranges.

Here, using cryo-electron microscopy (cryo-EM) ^27^, we resolved the 3D structure of Alka and found that it assembles as a homopentamer with an extracellular domain and a transmembrane pore-forming region. Transitioning from near-neutral to alkaline pH induces conformational changes that shift the channel from a closed to a partially open state. Notably, we identify a unique lysine residue (K229) in the activation domain as a critical pH sensor. Deprotonation of its side chain under highly alkaline conditions disrupts stabilizing interactions, triggering structural rearrangements required for channel opening. Consistent with this, mutation of K229 markedly impairs high-pH activation of Alka in heterologous cells and diminishes alkaline taste sensation in flies. Together, our findings reveal how Alka senses and responds to highly alkaline pH, uncovering a distinct gating mechanism. This work defines a molecular principle of alkaline-pH detection through ion channels and provides broader insight into how ion channels diversify as chemosensory receptors to meet extreme chemical challenges.

## Results

### Alka cryo-EM structure at near-neutral pH

To elucidate the structural and functional mechanisms of the Alka channel in alkaline pH sensing, we set out to solve its 3D structure using single-particle cryo-EM. We first confirmed that Alka, when expressed in HEK293T cells, is activated by highly alkaline pH using whole-cell patch-clamp recordings. Next, we expressed and purified full-length Alka proteins in suspension-adapted HEK293 cells ^27–29^. After harvesting and lysing the cells, we extracted the membrane proteins using n-dodecyl-β-D-maltoside (DDM) and cholesteryl hemisuccinate (CHS), and purified Alka with Strep-Tactin affinity chromatography and size-exclusion chromatography (SEC) under nearly neutral pH condition (pH 7.4) ^27–29^. We then pooled and concentrated the peak fractions and assessed sample purity by SDS-PAGE followed by Coomassie blue staining. These analyses confirmed that the fly Alka proteins were highly pure, enabling us to prepare cryo-EM grids and examine their 3D structures. After two-dimensional (2D) classification and 3D reconstruction ^27,30^, a 3.5 angstrom (Å) cryo-EM density map was obtained. The high-resolution density maps facilitated accurate atomic model building for the majority of the protein, with clear visualization of side-chain densities and other fine structural features. This enabled high-resolution structural determination of Alka in its closed conformation. Our cryo-EM analysis of Alka revealed a homo-pentameric assembly adopting a barrel-stave architecture, measuring 86 Å in width and 112 Å in height (**Figure 1a**). Both the N- and C-termini of Alka are located on the extracellular side. Each Alka protomer mainly consists of an extracellular domain (ECD) and a transmembrane domain (TMD) (**Figure 1b**). The pentameric assembly of Alka features a central ion-conduction pore aligned along the fivefold axis of symmetry (**Figures 1c, 1d**). Specifically, the TMD consists of four α-helices (M1–M4) arranged around the central pore, with the M2 helices lining the pore axis and containing key residues essential for ion selectivity and gating (**Figures 1c, 1d**). The ECD of Alka is primarily composed of ten β-strands arranged in a β-sandwich topology, mediating interactions with neighboring subunits. Key loops, including Cys-loop, Loop A, Loop B, and Loop C, are positioned between these β-strands (**Figures 1e–g**). Notably, Loop C is uniquely situated at the interface between adjacent subunits, bridging protomer interactions within the homopentameric Alka channel complex (**Figures 1f, 1g**). This strategic placement makes Loop C a critical site for allosteric regulation ^31,32^. The homopentameric architecture of Alka bears structural resemblance to canonical pLGICs ^18^ such as glycine receptors (GlyRs) despite limited sequence similarity (30%) ^6,20,21^,reflecting a shared evolutionary origin and potentially similar gating principles.

**Figure 1.**
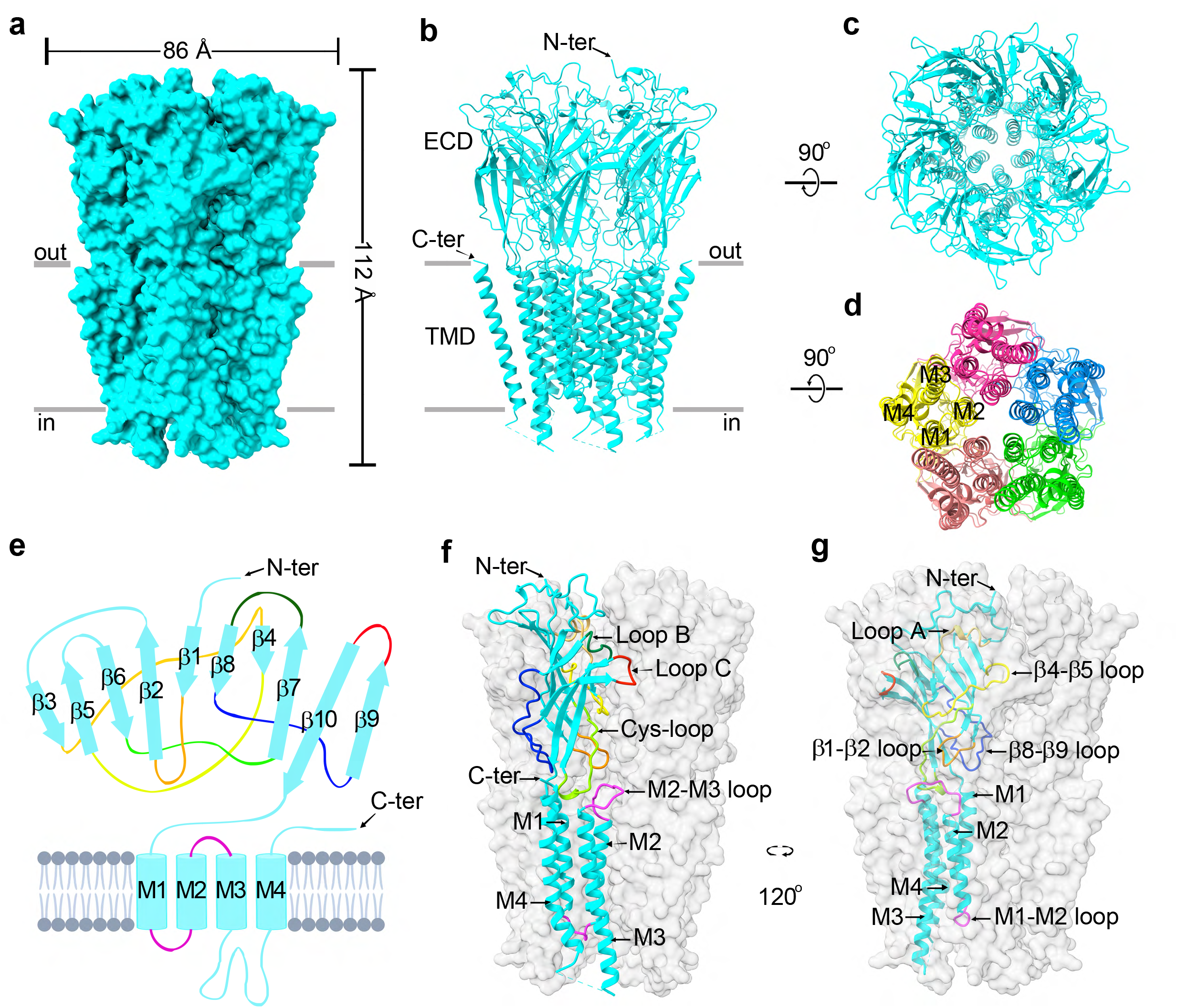
Cryo-EM structures of the Alka channel at pH 7.4. (**a**) Molecular surface of the Alka channel at pH 7.4, showing its homopentameric barrel-like architecture in side view. (**b**) Side view of the atomic model of the Alka channel at pH 7.5, highlighting the extracellular domain (ECD) and transmembrane domain (TMD). The amino terminus (N) and carboxyl terminus (C) are indicated. (**c, d**) Top (**c**) and bottom (**d**) views of the Alka channel structure, oriented perpendicular to the membrane. The channel consists of five identical subunits, each shown in a different color in (**d**), with the positions of the M1– M4 transmembrane segments labeled in (**d**). (**e**) Schematic diagram of the topology and secondary structure elements of the Alka channel, showing the N- and C-termini. (**f, g**) Structure of a single subunit highlighted in blue within the Alka channel complex. Loops are shown in different colors.

### Conformational changes of Alka at alkaline pH

To investigate how Alka responds to alkaline pH, we determined its structure under activating conditions. We incubated purified Alka protein in pH 10.0 buffer and rapidly vitrified the samples for cryo-EM analysis ^27^. We resolved the structure at pH 10.0, although protein aggregation prevented us from examining even higher pH. At pH 10.0, Alka assembles into a homopentamer, with five identical protomers arranged symmetrically around a central fivefold axis. The overall architecture at pH 10.0 remains broadly similar to that observed at near-neutral pH, consisting primarily of an ECD and a TMD. In the TMD, each protomer contains four α-helices (M1–M4), and the M2 helices from all subunits converge to form the ion conduction pore along the central axis (**Figures 2a–d**). Despite this general similarity, careful inspection of Alka’s 3D architecture reveals a shape change at pH 10.0 compared with pH 7.4. Notably, the Alka channel complex measures approximately 81 Å in diameter and 118 Å in height at pH 10.0 (**Figure 2a**), adopting a more compact and elongated conformation than at pH 7.4, which measures 86 Å in diameter and 112 Å in height (**Figure 1a**). The reduction in diameter reflects tighter packing of the subunits around the central axis, while the increase in height indicates elongation along the vertical axis. Both the ECD and the TMD contribute to these shape changes, suggesting that alkaline conditions prime the channel for ion conduction.

**Figure 2.**
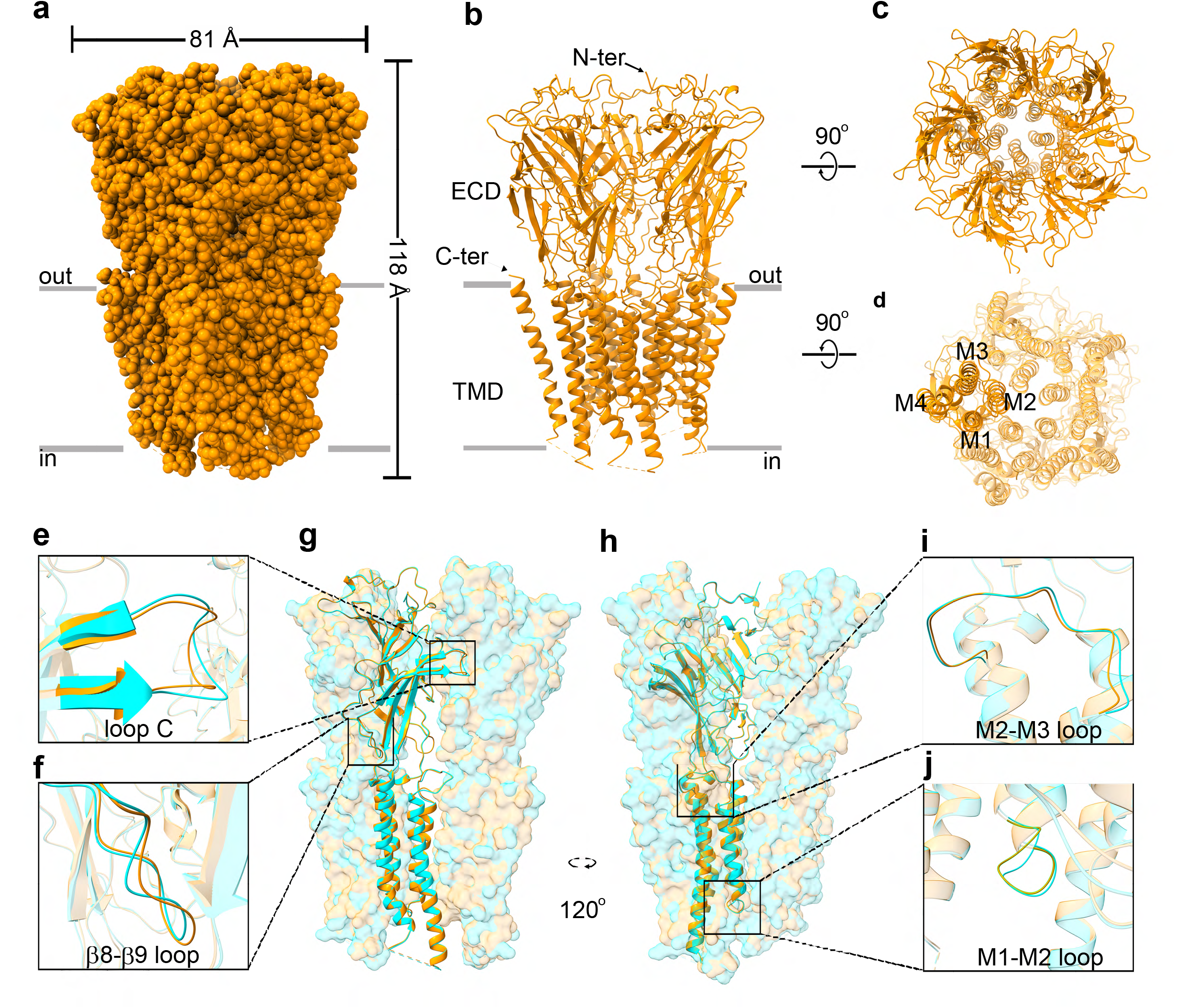
Cryo-EM structures of the Alka channel at pH 10.0. (**a**) 3D cryo-EM structure of Alka at pH 10.0, showing a more compact and elongated conformation compared to pH 7.4. (**b**) Side view of the homopentameric Alka structure at pH 10.0, highlighting the extracellular domain (ECD), transmembrane domain (TMD), and the N- and C-termini. (**c, d**) Top (**c**) and bottom (**d**) views of the Alka channel, showing its five-fold symmetry. Each subunit contains four transmembrane segments (M1–M4), with the pore-lining M2 helices forming the conduction pathway. (**e–i**) Conformational rearrangements of several extracellular loops between pH 7.4 and pH 10.0, including Loop C (**e**), β8–β9 loop (**f**), M1–M2 linker (**j**), and M2–M3 linker (**i**).

Notably, comparison of Alka structures at pH 7.4 and 10.0 revealed concerted conformational rearrangements that facilitate gating (**Figures 2e–g**). The most pronounced change occurs in Loop C, which undergoes an outward displacement and partial loss of helical structure, extending away from the extracellular core (**Figure 2e**).

This structural relaxation likely alleviates steric constraints and promotes flexibility required for activation. Additional but more subtle changes are observed in the β8–β9, M1–M2, and M2–M3 loops (**Figures 2f, 2i, 2j**), suggesting that pH sensing triggers a coordinated reorganization of both extracellular and transmembrane domains. Among these elements, Loop C shows the largest displacement, consistent with its established role in ligand gating and activation in pLGICs ^27,31,32^. We therefore propose that alkaline pH, or hydroxide ions (OH^-^), acts as a ligand that directly engages Loop C to induce structural rearrangements culminating in channel opening.

As observed in many pLGICs that function as neurotransmitter receptors, including glycine receptors (GlyRs), GABA type A receptors (GABAARs), and nicotinic acetylcholine receptors (nAChRs), Loop C contributes to the ligand-binding pocket for neurotransmitters, such as acetylcholine ^24^, glycine ^20,33,34^, and GABA ^22^. Upon ligand binding, Loop C undergoes conformational rearrangements that propagate across the ECD and are transmitted to the TMD, initiating channel opening. The ligand-binding pockets in GlyRs, GABAARs, and nAChRs are relatively wide, allowing accommodation of small molecule ligands. By contrast, in Alka, Loop C adopts a narrower, occluded conformation that sterically blocks access to the binding cavity. This structural feature aligns with our previous electrophysiological data showing that Alka does not respond to classical ligands such as GABA or Glycine ^6^, indicating a distinct activation mechanism.

Moreover, we compared the pore architecture of Alka at pH 7.4 (**Figure 3a**) and pH 10.0 (**Figure 3b**). In both states, the narrowest constriction resides at Pro276, a conserved residue within the pore-lining M2 helix ^6^ (**Figures 3a, 3b**). At pH 7.4, the minimal pore radius is 1.7 Å (**Figures 3a, 3c**), insufficient to accommodate Cl^−^ ions (1.8 Å radius), consistent with a closed, non-conducting conformation. In contrast, at pH 10.0, the pore modestly dilates to 1.9 Å (**Figures 3b, 3c**), reflecting a transition toward a partially open conformation capable of supporting limited Cl^−^ permeation. Alternatively, Alka could rapidly activate and then enter a desensitized state. These observations indicate that alkaline pH triggers conformational rearrangements that widen the pore, thereby facilitating ion conduction.

**Figure 3.**
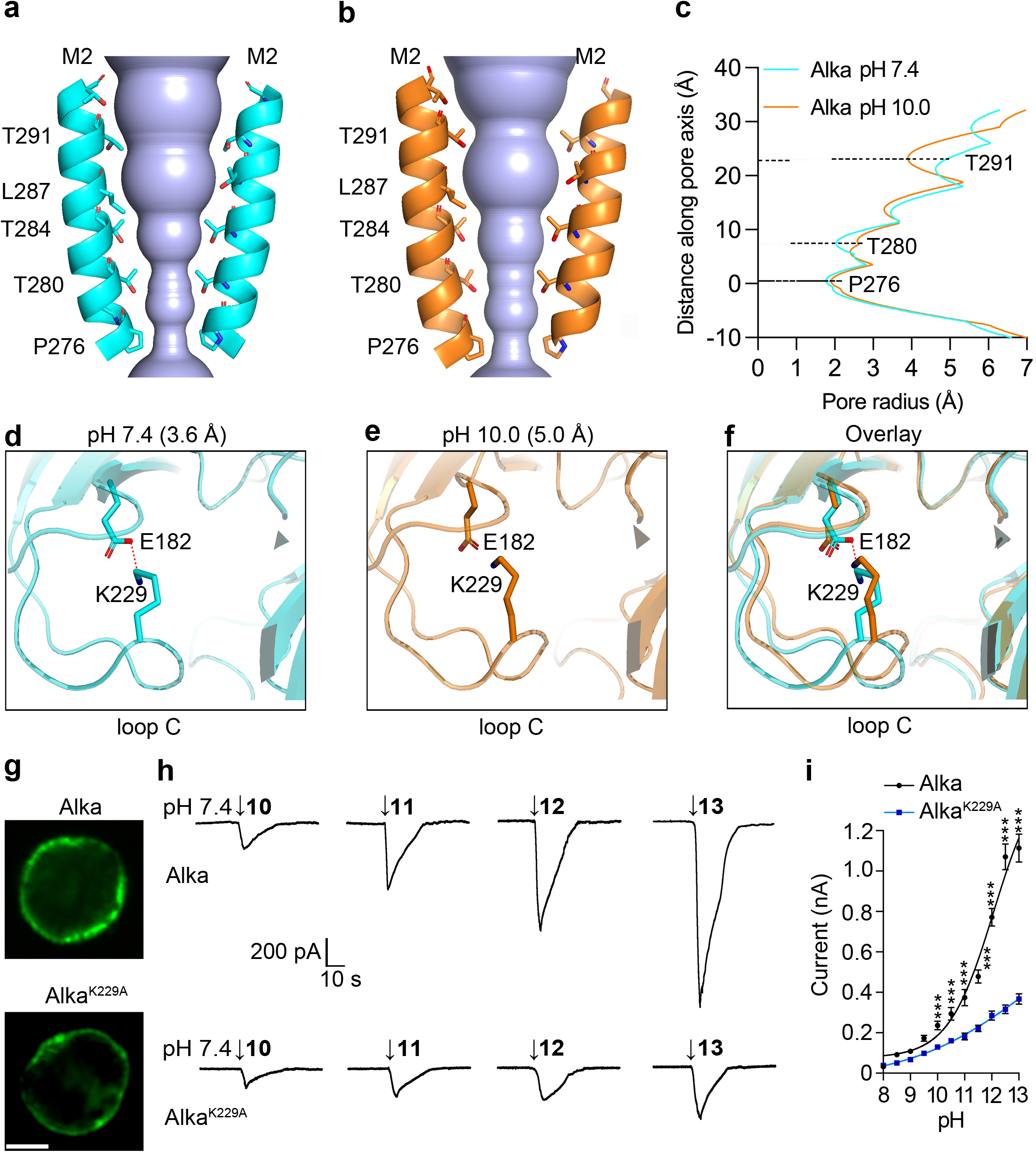
Alkaline pH triggers conformational changes in the Alka pore and Loop C domains. (**a, b**) Pore profiles of the Alka channel at pH 7.4 (blue) and pH 10.0 (brown). The M2 helices of two diagonal subunits are shown in cartoon representation, and pore-lining residues are depicted as sticks. (**c**) Comparison of the pore radius along the pore axis for Alka at pH 7.4 and pH 10.0. (**d**) Ligand-binding Loop C domain at pH 7.4, showing a salt bridge between K229 and E182 that stabilizes the closed conformation of the ligand-binding pocket. (**e**) Loop C at pH 10.0, with displacement of K229 relative to E182, resulting in disruption of the salt bridge and opening of the Loop C pocket. (**f**) Overlay of Loop C structures at pH 7.4 and 10.0, highlighting the K229 side chain movement associated with pH-dependent conformational changes. (**g**) Confocal images of HEK-293 cells expressing wild-type Alka or the K229A mutant. Scale bar, 5 µm. (**h**) Whole-cell currents evoked by alkaline solutions (pH 10–13) in HEK-293 cells expressing wild-type Alka or the K229A mutant. (**i**) Dose responses of inward currents to varying alkaline pH solutions for wild-type Alka and the K229A mutant. n=11 cells, \*\*\**p<*0.001, ANOVA tests.

### Role of K229 in alkaline pH sensing

Given that the inter-subunit positioning of Loop C is essential for coordinating the transition from the closed to the open state for the pGLIC superfamily ^27,31,32^, we propose that the Loop C region serves as a critical site for pH-dependent gating and activation of the Alka channel. Comparative sequence analysis reveals that fly Alka harbors a unique lysine residue at position 229 (K229) within Loop C, which is absent in GlyRs, GABAARs and nAChRs across species. Given that the pKa (~10.0) of the ε-amino group on lysine is close to Alka’s activation pH threshold, we hypothesize that K229 undergoes deprotonation in response to alkaline pH. This deprotonation could disrupt local electrostatic interactions or hydrogen bonds, allowing K229 to act as an intrinsic alkaline pH sensor. To test this, we examined Loop C conformation at pH 7.4 and pH 10.0. At pH 7.4, K229 forms a salt bridge with the negatively charged glutamate 182 (E182), stabilizing Loop C (**Figure 3d**). At pH 10.0, the distance between K229’s ε-amino group and E182’s carboxylate increases from 3.6 Å to 5.0 Å, consistent with lysine deprotonation (**Figure 3e**). This elongation greatly weakens or even abolishes the salt bridge, causing local destabilization and loosening of Loop C (**Figure 3f**). We postulate that these structural changes propagate through the channel, priming it for activation under alkaline conditions.

To determine the functional significance of this structural transition, we generated a point mutation at K229, replacing lysine with alanine to remove its positive side chain. Confocal imaging of HEK293 cells expressing either wild-type or K229A Alka channels showed that both proteins efficiently localized to the plasma membrane, allowing us to perform whole-cell patch-clamp recordings ^6^ to examine channel responses to alkaline solutions (pH 10.0–13.0) applied locally to the cell surface (**Figure 3g**). Our recordings revealed that cells expressing Alka K229A exhibited markedly reduced amplitudes of alkaline pH–evoked currents compared to cells expressing wild-type Alka (**Figures 3h, 3i**). Since the K229A mutation markedly reduces channel currents, our results suggest that deprotonation of K229 at high pH triggers conformational changes from Loop C toward the pore domain, leading to channel opening. Substituting K229 with alanine eliminates this energetic linkage and disrupts allosteric coupling, preventing the pore from opening.

To assess the functional significance of the K229 residue of Alka in vivo, we tested whether this residue was required for high-pH detection in intact flies. Using the UAS/Gal4 system ^35^, we expressed either wild-type Alka (*UAS-Alka*) or the K229A mutant Alka (*UAS-Alka*^*K229A*^) in Alka-expressing cells (alka-Gal4) within an *alka*^*1*^ null mutant background ^6^. As reported previously ^6^, loss of *alka* caused severe defects in both physiological and behavioral responses to alkaline stimuli (pH 10.0–13.0). We examined whether expression of wild-type or K229A mutant Alka could restore these responses. Our tip recordings from gustatory receptor neurons (GRNs) ^5,6^ showed that expression of wild-type Alka generated robust trains of spikes in response to highly alkaline pH (10.0-13.0). In contrast, GRNs expressing the K229A mutant displayed significantly fewer spikes, indicating that the K229A channel was severely impaired in mediating high-pH activation (**Figures 4a, 4b**). Therefore, our results demonstrated that K229 was essential for Alka-dependent neuronal activation by alkaline stimuli. To determine whether these physiological defects translated into altered feeding behavior, we next performed two-choice feeding assays ^5,6,36-39^. We found that wild-type flies consistently avoided alkaline food when presented alongside neutral food. Expression of wild-type Alka in the *alka*^*1*^ mutant background restored this avoidance behavior, whereas animals expressing the K229A mutant failed to discriminate highly alkaline food from neutral food (**Figure 4c**). We further analyzed feeding responses using the proboscis extension reflex (PER) assay, in which we directly applied neutral or alkaline food to the proboscis ^5,6^. Wild-type flies strongly rejected alkaline food, showing a marked reduction in PERs. Expression of wild-type Alka in *alka*^*1*^ mutants restored this response, whereas expression of the K229A mutant did not (**Figure 4d**). Together, our results demonstrated that the K229 residue was critical for Alka-mediated taste responses to highly alkaline food in flies.

**Figure 4.**
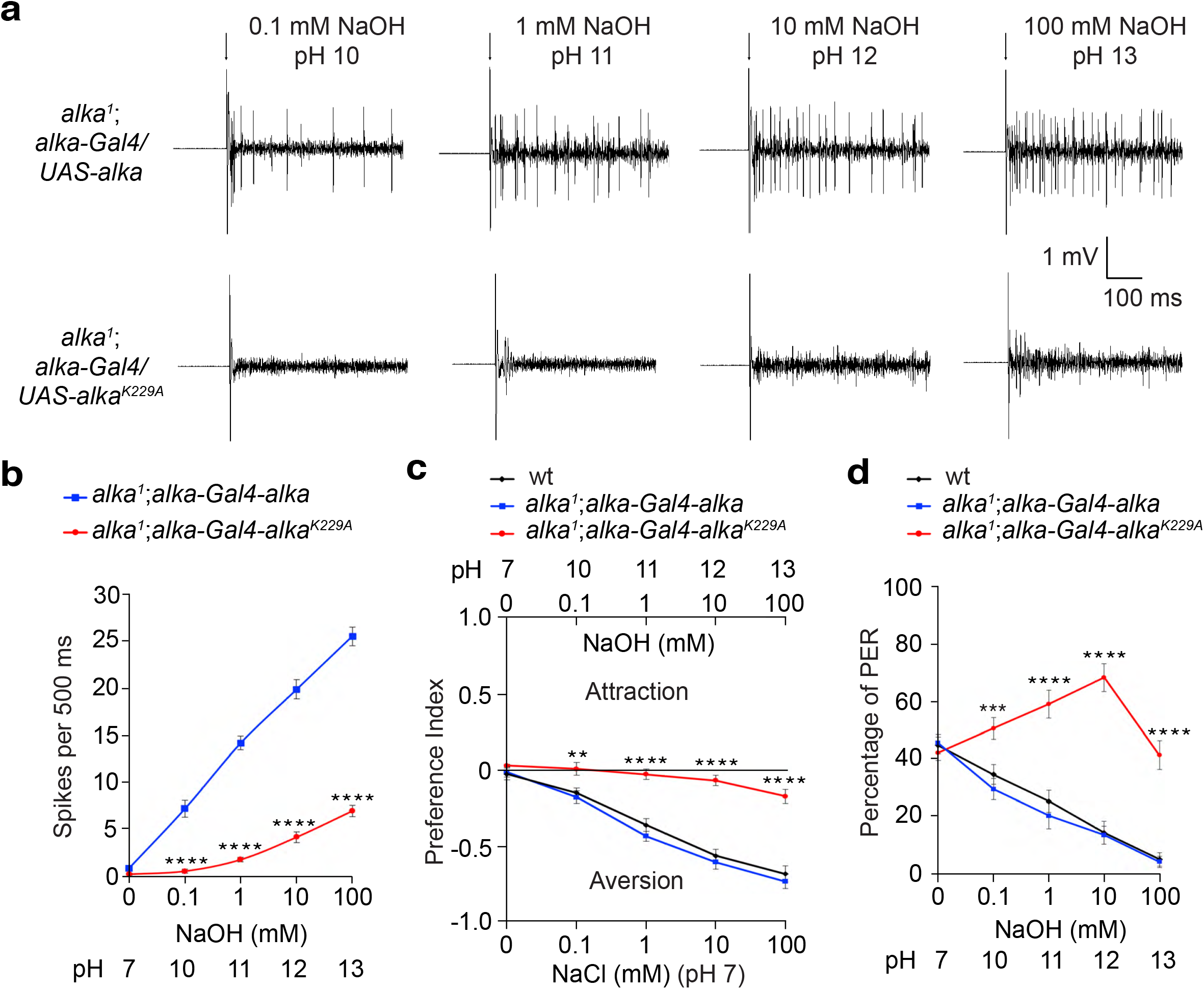
Lysine 229 (K229) in Loop C is critical for taste responses to alkaline pH. (**a**) Neuronal spikes recorded from gustatory receptor neurons (GRNs) in response to alkaline solutions (pH 10–13) in flies expressing wild-type Alka or the K229A mutant in an *alka*^*1*^ null mutant background. (**b**) Quantification of spike frequency in flies expressing wild-type Alka or the K229A mutant in an *alka*^*1*^ null mutant background. n = 12 animals. \*\*\*\**p* < 0.0001, ANOVA tests. (**c**) Two-way feeding assays showing preference between neutral food (NaCl) and alkaline food (NaOH) in wild-type flies and *alka*^*1*^ mutants expressing wild-type Alka or Alka K229A. n = 12 trials. \*\*\**p* < 0.001, ANOVA. (**d**) Proboscis extension response (PER) assays to liquid food containing 50 mM sucrose with varying concentrations of NaOH in wild-type flies and *alka*^*1*^ mutants expressing wild-type Alka or Alka K229A. n = 12 trials. \*\*\*\**p* < 0.0001, ANOVA tests.

In conclusion, we uncover a pH-dependent gating mechanism for Alka, in which alkaline conditions initiate conformational changes centered on Loop C. At near-neutral pH, the channel remains in a closed conformation, stabilized by a salt bridge between K229 in Loop C and E182, thereby maintaining a non-conductive state. Under highly alkaline conditions, deprotonation of K229 disrupts this electrostatic interaction, driving a closed-to-partial-open transition and enabling chloride flux. This mechanism identifies alkaline pH or OH^-^ as the activating stimulus, defining Alka as a channel specialized for alkaline pH sensing rather than classical ligand binding. By revealing how a single lysine residue has been uniquely adapted as a molecular pH sensor, our findings provide a framework for understanding how ion channels diversify to detect extreme environmental conditions. Although the lysine-based sensor (K229) appears unique to *Drosophila*, the principle that hydroxide ions function as direct gating ligands expands the known strategies of ion channel activation. This sensory receptor–level innovation underscores the adaptability of living organisms in devising new molecular solutions to environmental challenges. More broadly, our work highlights molecular principles of alkaline pH detection that may be independently adopted across other organisms, offering new perspectives on the evolutionary plasticity of sensory systems ^40^.

## Author contributions

T.M., Y.W., Y.Z., and Y.V.Z. conceived and designed the experiments. Y.W. and T.M. expressed and purified Alka proteins. Y.W. and T.M. collected and analyzed the Cryo-EM data. T.M. performed whole-cell patch clamp recordings. Y.V.Z. performed tip recordings and feeding behavioral assays in flies. T.M., Y.W., Y.Z., and Y.V.Z. analyzed the data, prepared the figures, and wrote the manuscript.

## Competing interests

The authors declare that they have no competing interests related to this study.

## Data availability

All data supporting the findings of this study are included in the manuscript, main figures, and supplementary figures and materials. Additional information is available upon request from Y.Z. and Y.V.Z.

## Notes

### Competing Interest Statement

The authors have declared no competing interest.

